# STK25 directly activates LATS1/2 independent of MST/MAP4Ks

**DOI:** 10.1101/354233

**Authors:** Sanghee Lim, Nicole Hermance, Tenny Mudianto, Hatim M. Mustaly, Ian Paolo Morelos Mauricio, Marc A. Vittoria, Ryan J. Quinton, Brian W. Howell, Hauke Cornils, Amity L. Manning, Neil J. Ganem

## Abstract

The Hippo pathway maintains tissue homeostasis by negatively regulating the oncogenic transcriptional co-activators YAP and TAZ. Though functional inactivation of the Hippo pathway is common in tumors, mutations in core pathway components are rare. Thus, understanding how tumor cells inactivate Hippo signaling remains a key unresolved question. Here, we identify the kinase STK25 as a novel activator of Hippo signaling. We demonstrate that loss of STK25 promotes YAP/TAZ activation and enhanced cellular proliferation, even under normally growth-suppressive conditions. We reveal that STK25 activates LATS via a previously unobserved mechanism, in which STK25 directly phosphorylates the LATS activation loop. This represents a new paradigm in Hippo activation and distinguishes STK25 from all other identified kinase activators of LATS. *STK25* is significantly focally deleted across a wide spectrum of human cancers, suggesting *STK25* loss may represent a common mechanism by which tumor cells functionally impair the Hippo tumor suppressor pathway.

---

First discovered in *Drosophila* as a critical regulator of organ size, the Hippo tumor suppressor pathway has emerged as playing a major role in maintaining tissue homeostasis through the regulation of cell proliferation and survival (Zanconato et al. 2016). The key mediators of Hippo signaling are LATS1 and LATS2 (Large Tumor Suppressor) kinases, which function to negatively regulate the activity of the oncogenic transcriptional co-activators Yes-associated protein (YAP) and transcriptional co-activator with PDZ-binding motif (TAZ) (Zhang et al. 2009, Zhao et al. 2010). Upon stimulation of Hippo signaling, activated LATS kinases directly phosphorylate YAP/TAZ at conserved serine residues, which promote YAP/TAZ nuclear extrusion and subsequent degradation (Zhao et al. 2010). By contrast, in the absence of LATS activation, YAP/TAZ are free to translocate into the nucleus, where they bind to the TEAD/TEF family of transcription factors to promote the expression of numerous genes essential for cellular proliferation and survival (Wu et al. 2008, Zhang et al. 2008, Hong et al. 2005). Deregulation of LATS1/2 kinases, which leads to subsequent hyper-activation of YAP/TAZ, is sufficient to promote tumorigenesis in mouse models (Zhou et al. 2009, Nishio et al. 2015). Furthermore, amplification of YAP and/or TAZ has been found in a variety of human malignancies (Overholtzer et al. 2006, Fernandez-L et al. 2009).

Multiple signals lead to the activation of LATS kinases, including contact inhibition, cellular detachment, loss of actin cytoskeletal tension, serum deprivation, glucose starvation, signaling from G-protein coupled receptors, and cytokinesis failure (Zhao et al. 2010, Dupont et al. 2009, Adler et al. 2013, Mo et al. 2015, Wang et al. 2015, Ganem et al. 2014, Yu et al. 2012, Dutta et al. 2018). Mechanistically, LATS kinases were initially found to be regulated by MST1 and MST2, the mammalian orthologs of the Drosophila Hpo kinase, after which the pathway is named. Activation of LATS1/2 initiates with the recruitment of MST1/2 to LATS kinases via interactions with scaffolding proteins such as SAV1, MOB1, and NF2 at the plasma membrane (Zhang et al. 2010, Yin et al. 2012). Once recruited, MST1/2 phosphorylate LATS1/2 at their hydrophobic motifs to remove the auto-inhibitory conformations of LATS1/2, thereby allowing auto- and trans-phosphorylation interactions to take place at the activation loop motifs of LATS1/2. It is this phosphorylation at the activation loop that subsequently leads to full LATS kinase activity (Ni et al. 2015, Hoa et al. 2016). However, it has become increasingly clear that kinases which regulate the activation of LATS are not limited to MST1/2 in mammalian cells. Genetic deletion of MST1/2 fails to prevent full LATS activation, and YAP/TAZ phosphorylation remains intact in mice lacking MST1/2 (Zhou et al. 2009, Meng et al. 2015). Moreover, several conditions known to stimulate LATS activation, such as contact inhibition, serum starvation, and cell detachment, do so in a MST1/2-independent manner, suggesting evolutionary divergence from *Drosophila* in mammalian cells, as well as the presence of additional upstream kinases that control LATS activation (Ganem et al. 2014, Zhou et al. 2009, Dutta et al. 2018, Plouffe et al. 2016). Indeed, recent work has shown the presence of additional upstream kinases that control LATS activation outside of MST1/2, as several members of the MAP4K family have been identified as having overlapping roles in directly phosphorylating the hydrophobic motif of LATS kinases (Zheng et al. 2015, Meng et al. 2015). However, cells in which *MST1/2* and all *MAP4K*s have been collectively deleted with CRISPR-mediated approaches still induce LATS and YAP phosphorylation upon stimulation, albeit at significantly reduced levels, indicating that more upstream activators of LATS kinases exist (Meng et al. 2015, Plouffe et al. 2016). Given that Hippo pathway inactivation has been found across numerous tumor types, but mutations and deletions of core Hippo signaling components are rare, the identification of novel upstream activators of Hippo signaling carries the potential to uncover previously unappreciated tumor suppressor genes (Pan D 2010, Kelliher and O’Sullivan 2013).

To identify novel upstream kinases that regulate LATS activity, we performed a focused RNAi screen to identify kinases that contribute to LATS phosphorylation and subsequent YAP phosphorylation. This approach identified STK25 as a novel upstream activator of the LATS kinases, whose loss significantly promotes YAP/TAZ activity. Mechanistically, we demonstrate that STK25 phosphorylates LATS at the activation loop motif in the absence of hydrophobic motif phosphorylation, which distinguishes it from all of the other known LATS activating kinases discovered to date.

## Results

### A focused kinome screen identifies STK25 as a novel regulator of LATS-YAP phosphorylation

We performed a focused RNAi screen to identify kinases that are necessary for inducing YAP phosphorylation under conditions of activated Hippo signaling in the non-transformed IMR90 fibroblast cell line. We focused our kinome screen on members of the Sterile20 superfamily of kinases (which includes MST1/2), as members of this superfamily maintain structural similarities in spite of evolutionary divergence (Thompson and Sahai 2014). We depleted individual kinases via RNAi and then stimulated Hippo pathway activation by treatment with the drug dihydrocytochalasin B (DCB), which destabilizes the actin cytoskeleton and mimics activation of Hippo signaling under loss of F-actin driven cytoskeletal tension (MacLean-Fletcher and Pollard 1980, Zhao et al. 2012). Treatment with DCB induced robust activation of Hippo signaling with consequent phosphorylation of YAP at serine 127, which is a LATS-specific canonical site that regulates YAP cytoplasmic retention through binding to 14-3-3 proteins (Zhao et al. 2010). As expected, we found that depletion of known activators of LATS, such as MST1/2, decreased YAP phosphorylation, but surprisingly we found STK25 to be the strongest hit in our screen (Supplementary Fig. 1a-d). Depletion of STK25 significantly reduced levels of YAP S127 phosphorylation as assessed by quantitative immunoblotting relative to controls in this assay (Supplementary Fig. 1a-d). Importantly, this effect was reproduced in multiple cell lines, including both HEK293A and hTERT RPE-1 (Fig. 1a, Supplementary Fig. 1c, d). We performed phos-tag gel electrophoresis to assess overall levels of YAP phosphorylation and found that STK25-depletion led to significantly reduced levels of phosphorylated YAP, with consequent enrichment of unphosphorylated YAP, as compared to controls (Fig. 1b). Additionally, we found that TAZ, a mammalian paralog of YAP, was also enriched in an unphosphorylated status in STK25-depleted cells (Fig. 1c). We reproduced these results in cells treated with Latrunculin A, which is a fungal-derived actin-binding toxin that has a different mechanism of action than cytochalasin-class agents (Supplementary Fig. 1e) (Morton et al. 2000).

**Figure 1.**
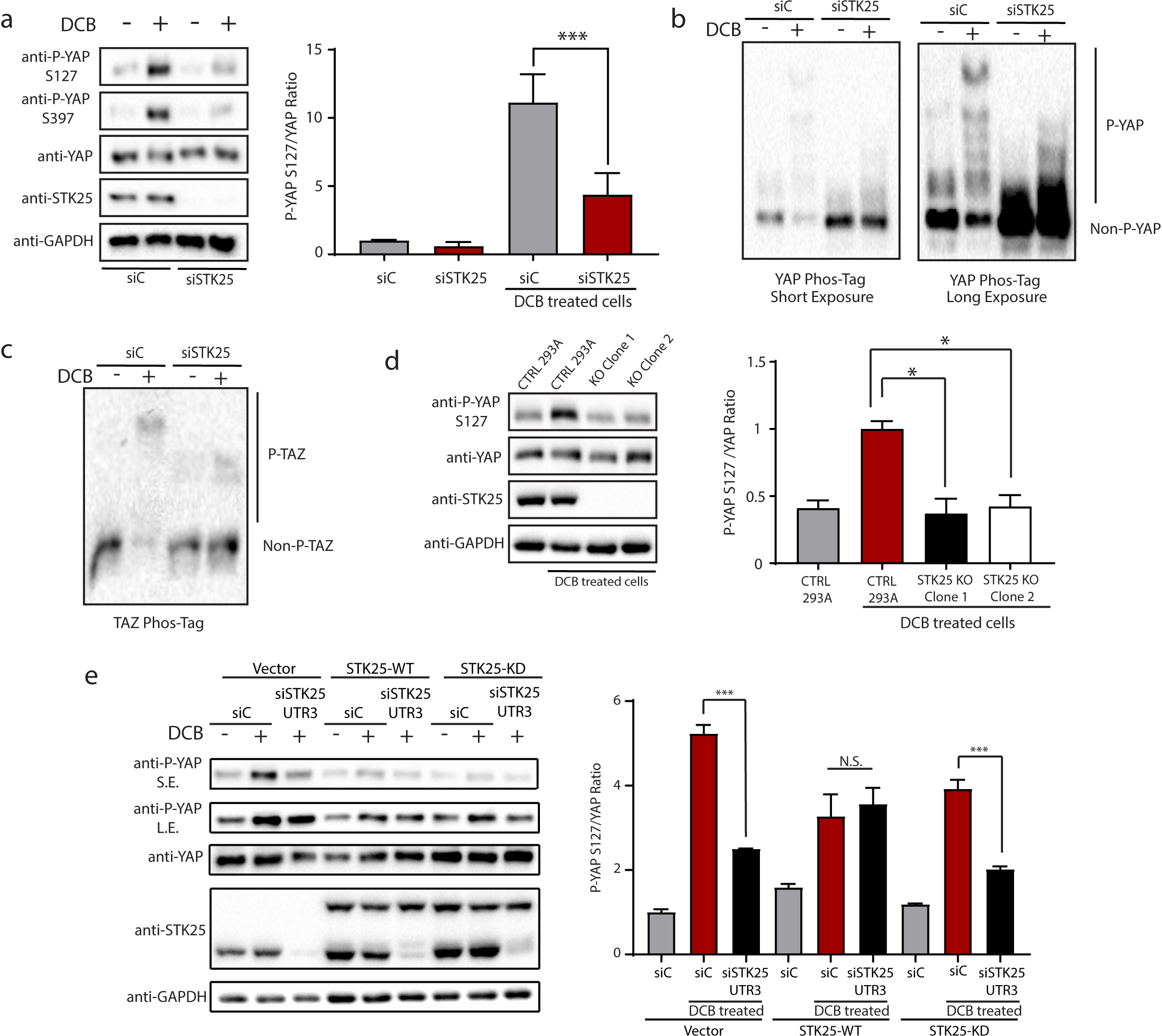
STK25 regulates Hippo activation in response to loss of cytoskeletal tension. **a.** Immunoblot and quantitation of phosphorylated YAP levels following treatment with 10 µM DCB in HEK293A cells transfected with the indicated siRNA. (n=6; ***p<0.001, unpaired t-test). **b**. Global phosphorylation status of YAP was assessed using Phos-tag gel electrophoresis following treatment with 10 µM DCB in HEK293A cells transfected with either control siRNA or STK25 siRNA. Shifted bands indicate degrees of YAP phosphorylation. **c.** TAZ phosphorylation status was assessed using Phos-tag gel electrophoresis following treatment with 10 µM DCB in HEK293A cells transfected with either control siRNA or STK25 siRNA. **d.** Immunoblot and quantitation of phosphorylated YAP levels following treatment with 10 µM DCB in either control HEK293A stably expressing Cas9 and a non-targeting sgRNA or STK25 KO 293A stably expressing Cas9 together with either sgRNA 1 (Clone 1) or sgRNA 2 (Clone 2) targeting STK25. (n=4; *p<0.05, One-way ANOVA with Dunnett’s post-hoc analysis). **e.** Immunoblot and quantitation of phosphorylated YAP levels following treatment with 10 µM DCB in HEK293A cells stably expressing wild-type STK25 (STK25-WT), kinase-dead STK25 (STK25-KD), or vector control (Vector) transfected with the indicated siRNAs. (n=3; ***p<0.001, N.S.: not significant; One-way ANOVA with Tukey’s post-hoc analysis). S.E. stands for Short Exposure; L.E. stands for Long Exposure.

We used multiple approaches to ensure that decreases in YAP phosphorylation following STK25 depletion via RNAi was not due to RNAi-induced off-target effects. First, we validated this finding with multiple distinct siRNA sequences targeting STK25 (8 total siRNAs, including three targeting the 3’UTR), and we observed that the degree of reduction in YAP phosphorylation strongly correlated with STK25 knockdown efficiency (Supplementary Fig. 1f, g). Second, we used CRISPR-Cas9 to genetically knockout STK25 from HEK293A cells and found that STK25 KO clonal cells (generated with two different sgRNA sequences) similarly failed to induce YAP phosphorylation to the same extent as control cells expressing Cas9 and a non-targeting sgRNA following DCB treatment (Fig. 1d). Finally, we demonstrated that overexpression of siRNA-resistant wild-type STK25, but not kinase-dead STK25 ^K49R^, was sufficient to rescue the loss of YAP phosphorylation observed in our knockdown experiments (Fig. 1e). Altogether, these data reveal that the kinase STK25 plays a previously unappreciated role in promoting YAP phosphorylation.

### STK25 Depletion Promotes YAP Activation

We next analyzed if the decrease in YAP phosphorylation following STK25 depletion leads to a corresponding increase in the nuclear localization of active YAP. Depletion of STK25, either by RNAi-mediated knockdown or CRISPR-mediated gene knockout, led to a significant increase in nuclear YAP in multiple cell lines (Fig. 2a-f, Supplementary Fig. 2d-i). We also observed a gene dose-dependent increase in nuclear YAP in *STK25^-/-^* and *STK25^+/-^* MEFs (Supplementary Fig. 2a-c). Remarkably, we found that depletion of STK25 enabled a population of YAP to remain nuclear even under conditions of actin depolymerization, which strongly sequesters YAP in the cytoplasm in control cells (Fig. 2a-c).

**Figure 2.**
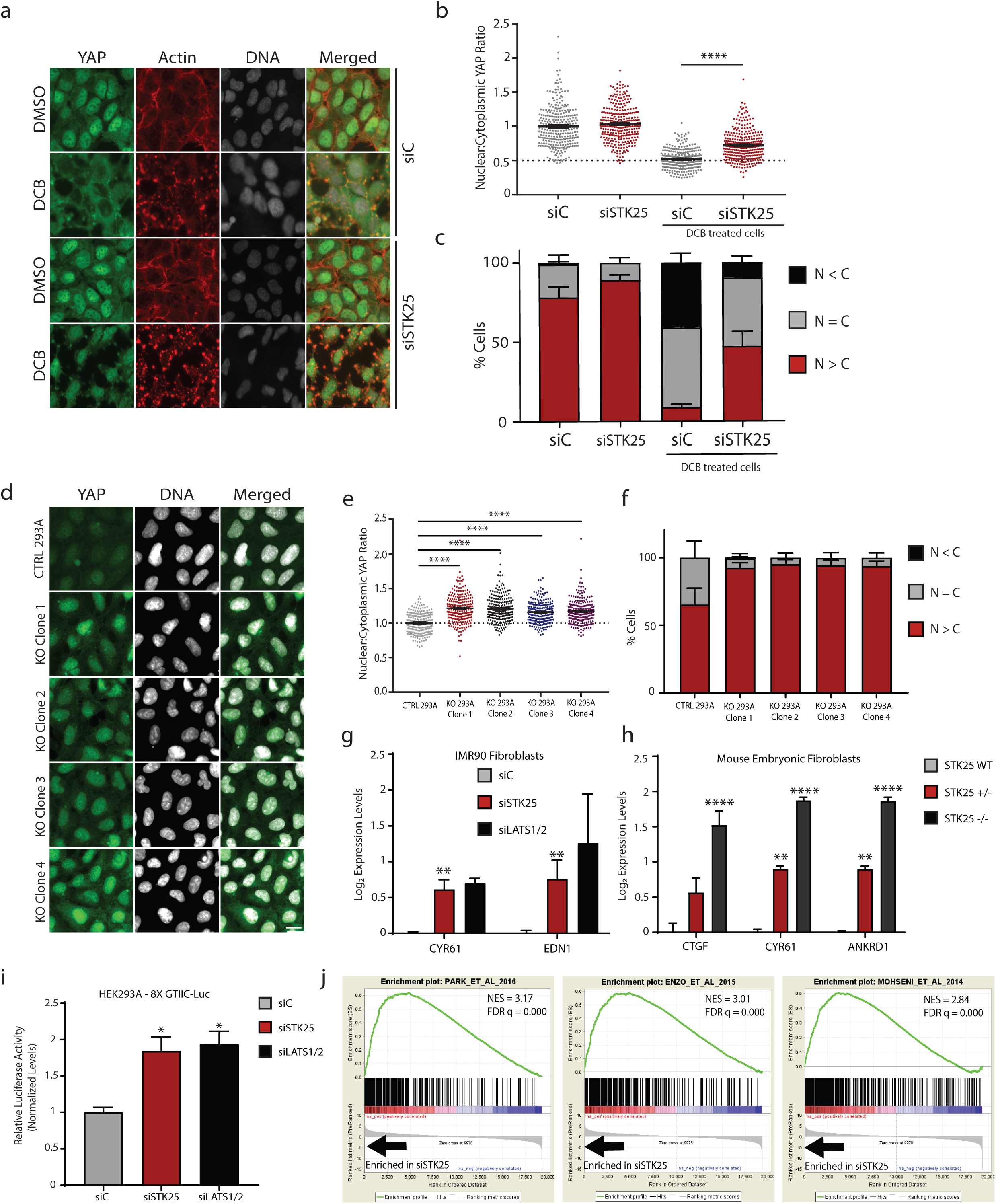
Loss of STK25 promotes activation of YAP. **a.** HEK293A cells transfected with either control siRNA or STK25 siRNA stained for YAP (Green), Actin (Red), and DNA (White) following treatment with 5 µM DCB. Scale bar, 20 µm. **b.** YAP intensity was quantified and nuclear:cytoplasmic ratios were calculated (n=225 per group over 3 biological replicates; ****p<0.0001, Mann-Whitney test). **c.**YAP localization was quantified (n=3; N>C, YAP is enriched in the nucleus; N=C, YAP is evenly distributed between the nucleus and the cytoplasm; N<C, YAP is enriched in the cytoplasm). **d.** Control and STK25 KO HEK293A were stained for YAP (Green) and DNA (White). Scale bar, 20 µm. **e.** Nuclear:cytoplasmic YAP ratios of control and STK25 KO HEK293A were quantified (n=225 per group over 3 biological replicates; ****p<0.0001, Kruskal-Wallis test). **f.** YAP localization in control and STK25 KO HEK293A was quantified as before (n=3 biological replicates). **g.** qPCR analysis of YAP-target gene expression in IMR90 fibroblasts transfected with the indicated siRNA (n=4; **p<0.01, unpaired t-test). **h.** qPCR analysis of YAP-target gene expression in wild-type, STK25^+/-^, and STK25^-/-^ mouse embryonic fibroblasts (n=3; **p<0.01, ****p<0.0001, One-way ANOVA with Dunnett’s post-hoc analysis). **i.** Expression of the TEAD luciferase reporter in HEK293A cells transfected with the indicated siRNA. Cells were transfected with siRNA, followed by transfection with 8X GTIIC TEAD luciferase reporter and pRL-TK renilla luciferase. Reporter luciferase activity was normalized to Renilla luciferase (n=3; *p<0.05, Oneway ANOVA with Dunnett’s post-hoc analysis). **j.** An expression signature of genes most upregulated upon loss of STK25 was constructed and GSEA was performed against a curated list of publicly available active YAP/TAZ gene sets. The top three most enriched gene sets are shown here.

To confirm that nuclear localized YAP was active, we first assessed whether depletion of STK25 increased YAP/TAZ activity using a luciferase-based gene expression reporter assay. HEK293A cells were transfected with a reporter encoding a YAP/TAZ-responsive luciferase gene, in which eight TEAD-YAP binding sites are cloned into the promoter driving expression of firefly luciferase (Dupont et al. 2011). Using this approach, we found that depletion of STK25 resulted in a doubling of expression from the luciferase reporter, indicating that loss of STK25 promotes YAP/TAZ activity (Fig. 2i). We also assessed the expression of YAP-target genes in cells depleted of STK25 and found that canonical YAP-target genes were significantly upregulated in IMR90 fibroblasts (Fig. 2g). Moreover, we found that YAP-target gene expression was also increased in a STK25 gene-dose dependent fashion in knockout MEFs for three well-established genes (Fig. 2h). Lastly, to assess gene expression in a comprehensive, unbiased fashion, we depleted STK25 from hTERT-RPE-1 cells and performed gene expression microarray analysis to obtain a list of genes that were significantly upregulated in cells lacking STK25 compared to controls. Gene Set Enrichment Analysis (GSEA) was performed using a curated set of 11 publically available, published gene sets for active YAP/TAZ as well as a composite gene set derived from RPE-1 cells expressing constitutively active YAP ^S5A^ or TAZ ^4SA^. This GSEA revealed that depletion of STK25 in RPE-1 cells results in a highly significant enrichment of active YAP/TAZ gene expression signatures (Fig. 2j, Supplementary Fig. 3a, b). Collectively, our data demonstrate that loss of STK25 promotes YAP/TAZ activation.

### STK25 acts through LATS1/2 to inhibit YAP

Given that loss of STK25 leads to an overall decrease in phosphorylation of YAP, we predicted that overexpression of STK25 may have the opposite effect and promote phosphorylation of YAP. Indeed, we found that overexpression of wild-type STK25, but not a kinase dead mutant (STK25 ^K49R^) led to significant increases in levels of phosphorylated YAP relative to controls (Supplementary Fig. 4a, b) and that this caused YAP to become enriched in the cytoplasm (Fig. 3a-c). We found that this effect of STK25 on YAP phosphorylation was LATS1/2-dependent, as overexpression of STK25 in HEK293A cells genetically depleted of LATS1/2 by CRISPR (LATS1/2 dKO) did not produce cytoplasmic enrichment of YAP (Fig. 3d-f). We also depleted STK25 from both wild-type and LATS dKO 293A cells and found that while loss of STK25 reduces YAP phosphorylation and drives YAP into the nucleus in wild-type cells, there was no such effect in LATS dKO 293A (Fig. 3g-i, Supplementary Fig. 4c), further indicating that STK25 functions through LATS1/2. Lastly, we found that knockdown of LATS1/2 via RNAi was sufficient to rescue YAP-target gene expression, even upon STK25 overexpression (Supplementary Fig. 4d). Thus, STK25 depends on LATS1/2 to regulate YAP.

**Figure 3.**
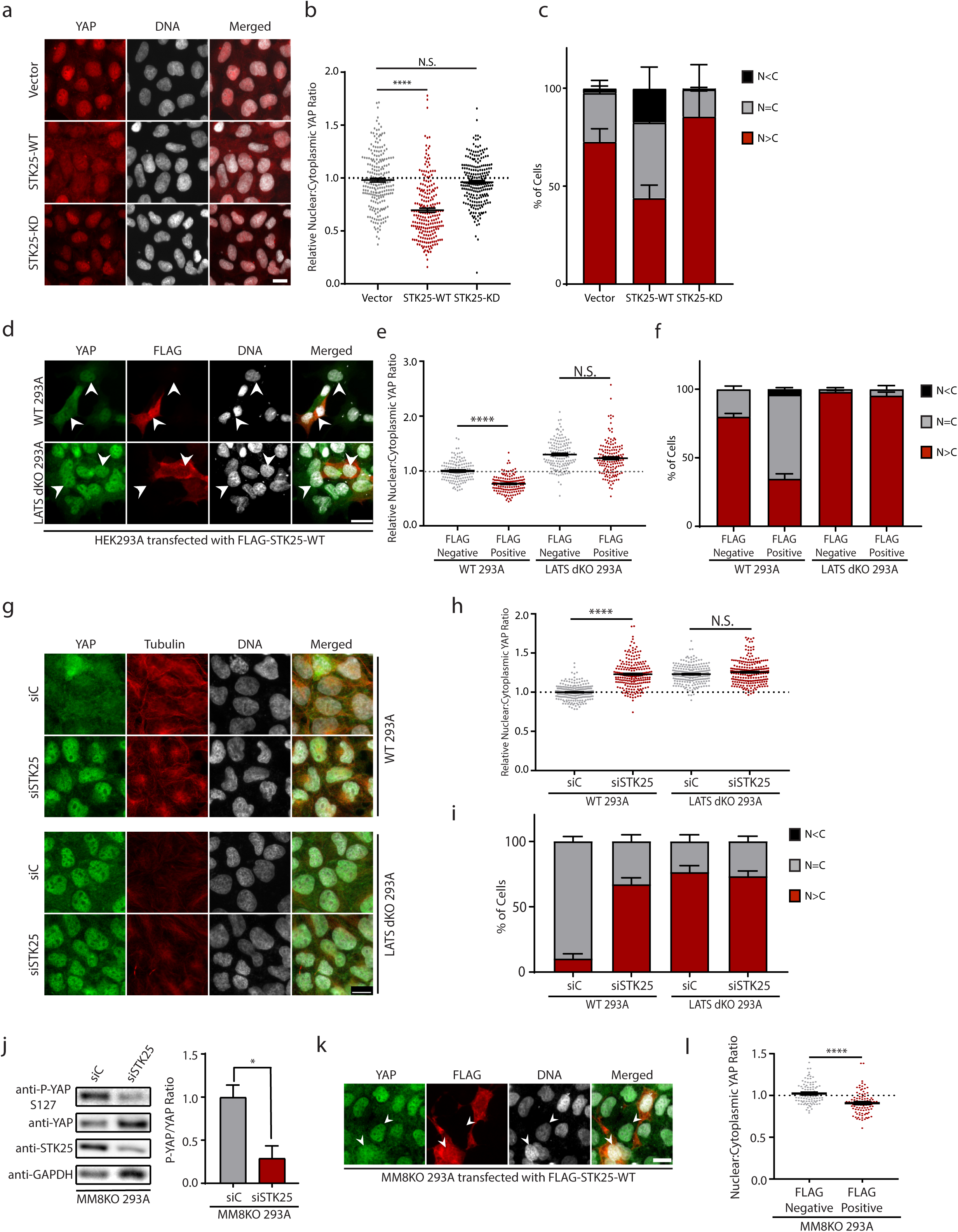
STK25 acts through LATS1/2 to inhibit YAP, independent of MST1/2 and MAP4Ks. **a.** HEK293A cells stably expressing STK25-WT, STK25-KD, or Vector were stained for YAP (Red) and DNA (White). **b.** YAP intensities from **a** were quantified and nuclear:cytoplasmic ratios were calculated (n=225 per group over 3 biological replicates; ****p<0.0001, Mann-Whitney test). **c.** YAP localization from **a** was quantified (n=3 biological replicates; N>C, YAP is enriched in the nucleus; N=C, YAP is evenly distributed between the nucleus and the cytoplasm; N<C, YAP is enriched in the cytoplasm). **d.** Wild-type and LATS dKO HEK293A were transfected with a vector encoding FLAG-STK25-WT and were stained for YAP (Green), FLAG (Red), and DNA (White). Arrows indicate representative cells selected for quantification that were positive for FLAG signal (indicating expression of transfected wild-type STK25) as well as an immediately adjacent cell negative of FLAG signal also selected for quantification to serve as controls. **e.** YAP intensities from **d** and nuclear:cytoplasmic ratios were calculated (n=200 per group over 4 biological replicates; ****p<0.0001, Kruskal-Wallis test with Dunn’s post-test; N.S. indicates “not significant”). **f.** YAP localization from **d** were quantified as before (n=4 biological replicates). **g.** Wild-type and LATS dKO HEK293A were transfected with the indicated siRNA, grown to confluence, then stained for YAP (Green), Tubulin (Red), and DNA (White). **h.** YAP intensities from **g** were quantified and nuclear:cytoplasmic ratios were calculated (n=225 over 3 biological replicates; ****p<0.0001, Kruskal-Wallis test with Dunn’s post-test; N.S. indicates “not significant.”) **i.** YAP localization from **g** were quantified as before (n=3 biological replicates). **j.** Immunoblot and quantification of phosphorylated YAP in MM8KO 293A cells transfected with the indicated siRNA (n=3; *p<0.05, paired t-test). **k.** MM8KO 293A cells were transfected with a vector encoding FLAG-tagged wild-type STK25 and were stained for YAP (Green), FLAG (Red), and DNA (White). **l.** YAP intensities from **k** were quantified in a FLAG-signal positive cell as well as an immediately adjacent FLAG-signal negative cell, and nuclear:cytoplasmic ratios were calculated (n=100 per group over 3 biological replicates; ****p<0.0001, Mann-Whitney test). All scale bars 20 µm, unless indicated otherwise.

Interestingly, this STK25-LATS1/2 axis is independent of other identified upstream LATS-activating kinases. HEK293A cells depleted of MST1, MST2, and all members of the MAP4K family (MAP4K1-7) via CRISPR-Cas9 (henceforth called MM8KO 293A) still demonstrate YAP phosphorylation when grown to confluence, albeit at much lower levels than in wild-type 293A. However, MM8KO 293A cells depleted of STK25 showed significantly decreased levels of YAP phosphorylation (Fig. 3j). Additionally, overexpression of STK25 in MM8KO 293A was sufficient to induce a cytoplasmic shift of YAP (Fig. 3k, l). This suggests that STK25 acts to regulate YAP in a LATS1/2-dependent fashion and that this signaling is independent of MST1/2/MAP4Ks’ activities.

### STK25 stimulates LATS kinases by promoting phosphorylation of LATS on the activation loop motif

Since STK25 is sufficient to induce YAP phosphorylation in the absence of other upstream LATS-activating kinases, and given the structural similarities between STK25 and MST1/2, we tested if STK25 directly phosphorylates and activates LATS. We first assessed whether STK25 and LATS interact. We expressed HA-tagged LATS2 and FLAG-tagged STK25 in HEK293A cells and found that immunoprecipitated HA-tagged LATS2 co-precipitated FLAG-tagged STK25 (Fig. 4a). To assess whether LATS1 or LATS2 bind to endogenous STK25, we overexpressed either Myc-tagged LATS1 or HA-tagged LATS2, and then immunoprecipitated each respective LATS kinase. We found that endogenous STK25 co-precipitated with both LATS1 and LATS2, suggesting that STK25 interacts with both LATS kinases to promote their activation (Fig. 4b).

**Figure 4.**
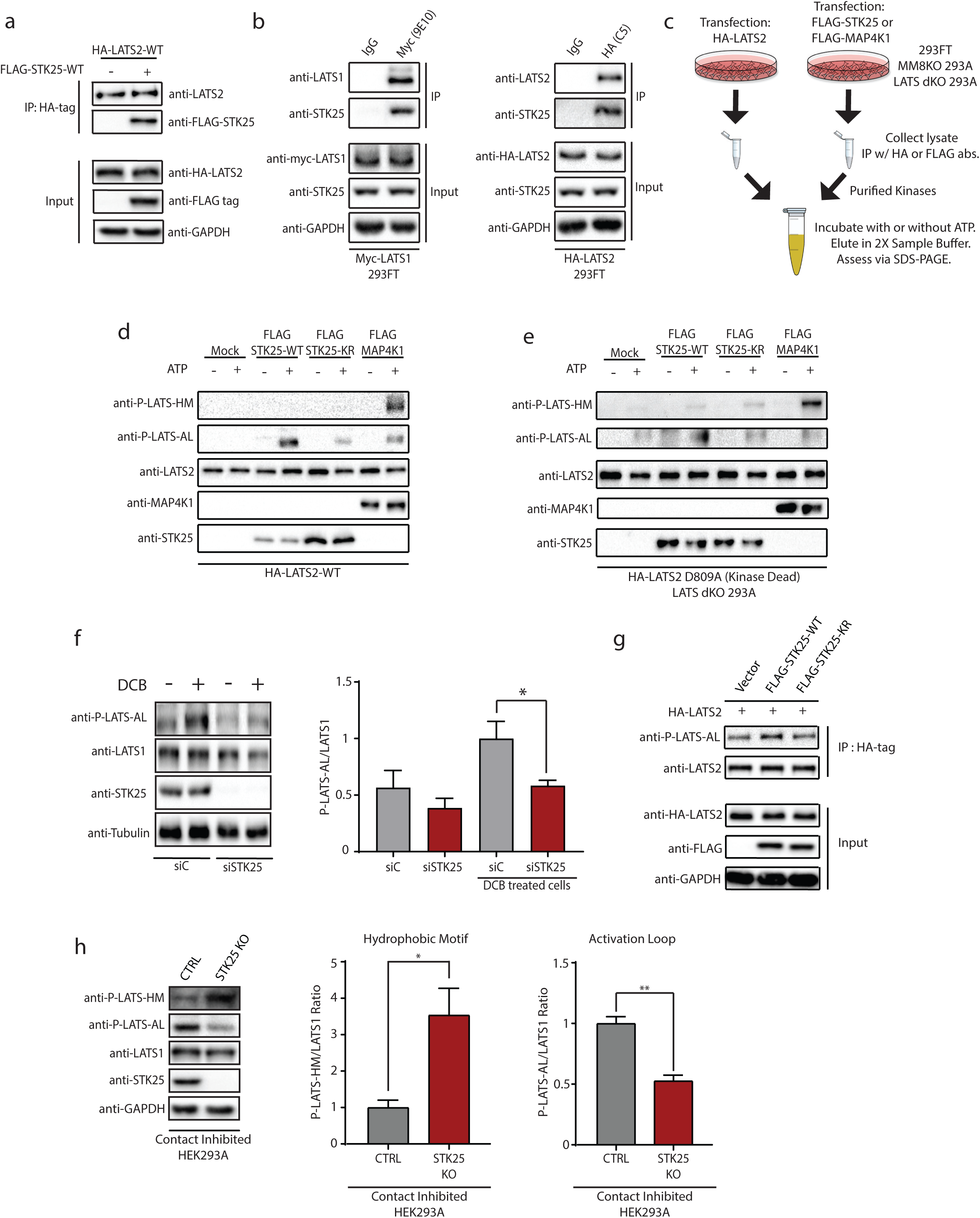
STK25 associates with LATS1/2 and directly phosphorylates LATS activation loop. **a.** LATS2 was immunoprecipitated from HEK293A cells co-transfected with HA-LATS2 and either vector control or FLAG-STK25. Co-precipitation of FLAG-STK25 with HA-LATS2 was assessed by immunoblotting. **b.** HEK293A cells were transfected with Myc-LATS1 or HALATS2. LATS1 and LATS2 were immunoprecipitated using antibodies directed against their tags and co-precipitation of endogenous STK25 was assessed by immunoblotting. **c.** Schema of the *in vitro* kinase assay set-up. The indicated cell lines were transfected with HA-LATS2, FLAG-STK25, or FLAG-MAP4K1. Protein lysates were then collected and used to individually immunoprecipitate kinases of interest. The purified kinases were then mixed together in kinase assay buffer and incubated for 30 minutes at 30°C with or without 500 µM ATP. Reactions were terminated by the addition of sample buffer, and levels of LATS phosphorylation were assessed via immunoblotting. **d.** Immunoprecipitation (IP) purified wild-type LATS2 (HA-LATS2-WT) was co-incubated with IP purified wild-type STK25 (FLAG-STK25-WT), kinase-dead STK25 (FLAG-STK25-KD), or wild-type MAP4K1 (FLAG-MAP4K1) and assessed for phosphorylation of its hydrophobic motif (P-LATS-HM) or activation loop (P-LATS-AL). **e.** IP purified kinase-dead LATS2 (HA-LATS2-KD) from transfected LATS dKO 293A cells was co-incubated with IP purified FLAG-STK25-WT, FLAG-STK25-KD, or FLAG-MAP4K1, all from transfected LATS dKO 293A. Levels of phosphorylated LATS at the hydrophobic motif (PLATS-HM) and activation loop (P-LATS-AL) were assessed via immunoblotting. **f.** Immunoblot and quantification of LATS activation loop phosphorylation (P-LATS-AL) following treatment with 10 µM DCB in HEK293A cells transfected with the indicated siRNA. (n=4; *p<0.05, paired t-test). **g.** HEK293A cells were co-transfected with HA-tagged wild-type LATS2 (HA-LATS2-WT) and either vector control (Vector), wild-type STK25 (FLAG-STK25-WT), or kinase-dead STK25 (FLAG-STK25-KD). LATS2 was immunoprecipitated and used to assess levels of activation loop phosphorylation by immunoblotting. Input lysates were assessed by immunoblotting for assessing protein loading and verification of transfected protein expression. **h.** Immunoblot and quantification of LATS1 hydrophobic motif (P-LATS-HM) and activation loop (P-LATS-AL) phosphorylation in either control HEK293A stably expressing Cas9 and a non-targeting sgRNA or STK25 KO 293A stably expressing Cas9 together with sgRNA 1 grown to confluence (n=3; *p<0.05, **p<0.01, paired t-test).

We next tested if STK25 could directly phosphorylate LATS. To do this, we carried out an *in vitro* kinase assay using HA-LATS2 as the substrate and either FLAG-STK25 or FLAG-MAP4K1 (positive control) as the kinase after proteins were immunoprecipitation-purified from transfected HEK293A lysates (Fig. 4c). As expected, we found that FLAG-MAP4K1 induced phosphorylation at the hydrophobic motif of LATS (LATS-HM), leading to subsequent phosphorylation at the activation loop (LATS-AL) (Fig. 4d). By contrast, STK25 did not induce phosphorylation at the LATS-HM, but was able to induce phosphorylation at the LATS-AL (Fig. 4d). This effect was dependent on STK25 kinase activity, as a kinase-dead STK25^K49R^ was unable to produce robust phosphorylation at the LATS-AL (Fig. 4d). Altogether, these data suggested that STK25 might be acting to activate LATS through direct phosphorylation at the activation loop site, bypassing phosphorylation at the hydrophobic motif.

To verify that STK25, and not other identified upstream activators of LATS kinases such as MST1/2 and members of the MAP4K family were responsible for the increase in phosphorylation at the activation loop of LATS, we utilized MM8KO 293A cells to IP purify STK25 and LATS2 for our *in vitro* kinase reactions as before. Once again, co-incubation of wild-type FLAG-STK25 with HA-LATS2 produced robust increases in phosphorylation at the activation loop of LATS2 as assessed by immunoblotting, which was decreased when LATS2 was co-incubated with kinase-dead STK25 ^K49R^ (Supplementary Fig. 5a). As phosphorylation at the activation loop has been canonically associated with LATS auto-phosphorylation, we wished to further verify that STK25 kinase activity, and not LATS2 intrinsic kinase activity, was driving this phenomenon (Tamaskovic et al. 2002, Stegert et al. 2004, Ni et al. 2015, Hoa et al. 2016). To accomplish this, we utilized LATS dKO 293A cells, into which kinase-dead LATS2 ^D809A^ was transfected. FLAG-STK25 and FLAG-MAP4K1 were also transfected into LATS dKO 293A cells. These proteins were IP-purified as before from the LATS dKO 293A cell lysates, which were then used for downstream *in vitro* kinase assays. Such a set-up ensured that we would be utilizing only LATS2 with no intrinsic kinase activity as our substrate, and that there would be no wild-type LATS1 or LATS2 that co-immunoprecipitated with either HA-LATS2 D809A, FLAG-STK25, or FLAG-MAP4K1 to confound interpretation of the data. Our results revealed that wild-type STK25, but not kinase-dead STK25 ^K49R^, promoted the phosphorylation of LATS-AL, independent of LATS intrinsic kinase activity (Fig. 4e). Further, we also noted that while MAP4K1 was able to promote phosphorylation of kinase-dead LATS2 at the hydrophobic motif, it was unable to do so at the activation loop. These data reveal that STK25 activates LATS kinases through a mechanism that is completely distinct from what has previously been characterized for the MST/MAP4K kinases (Fig. 4e).

To assess whether the increase in phosphorylated LATS-AL correlates with an increase in LATS activity, we transfected STK25 either alone, or together with LATS2, in LATS dKO 293A cells and then assessed levels of YAP phosphorylation via immunoblotting. As we previously noted, overexpression of STK25 in LATS dKO 293A was insufficient to induce YAP phosphorylation; however, transfection of wild-type LATS2 was sufficient to increase levels of phosphorylated YAP, which was further increased upon co-transfection of LATS2 with STK25 ^WT^, but not STK25 K49R (Supplementary Fig. 5b). Additionally, transfection of kinase-dead LATS2 abrogated this effect, which could not be rescued by co-transfection with STK25, strongly indicating once again that STK25 depends upon LATS kinases for its inhibitory effects on YAP (Supplementary Fig. 5b).

To further validate our *in vitro* findings that STK25 is able to phosphorylate LATS at its activation loop, we assessed the effects that modulation of STK25 has on LATS-AL phosphorylation in cells. To do this, we depleted STK25 via siRNA transfection and then stimulated LATS activity by treating cells with 10 µM DCB, after which the levels of activated LATS were assessed by quantitative immunoblotting. We observed an increase in phosphorylated LATS-AL with DCB treatment, which was significantly decreased upon knockdown of STK25 (Fig. 4f). Next, we overexpressed STK25 together with LATS2 in cells and found that co-expression of LATS2 along with wild-type STK25, but not kinase-dead STK25, was able to increase levels of phosphorylated LATS-AL (Fig. 4g). Lastly, we grew to confluence either control HEK293A cells expressing Cas9 together with a non-targeting sgRNA or STK25 KO HEK293A expressing Cas9 together with a sgRNA targeting STK25, and then assessed levels of phosphorylated LATS1 at the hydrophobic motif (LATS-HM) and the activation loop (LATS-AL) by quantitative immunoblotting. We found a significant decrease in phosphorylation of the LATS-AL in our STK25 KO HEK293A, but to our surprise, we also noted a robust and seemingly compensatory increase in phosphorylation of the LATS-HM (Fig. 4h). Further, this data suggested that cells lacking STK25 are unable to appropriately phosphorylate LATS-AL even when the LATS-HM is phosphorylated (Fig. 4h). Taken together, these data demonstrate that STK25 activates LATS through a previously unreported mechanism in which STK25 directly promotes phosphorylation of LATS-AL, independent from LATS intrinsic kinase activity.

### STK25 Loss Impairs LATS Activation under Physiologic Conditions and Provides a Proliferative Advantage

We next analyzed the effects of STK25 in physiologically relevant contexts that are known to stimulate LATS activation. We first assessed both LATS and YAP activity in confluent, contact inhibited cells, as contact inhibited cells are known to have strong activation of LATS kinases (Zhao et al. 2012). Indeed, we found that control HEK293A cells efficiently activated LATS and induced YAP phosphorylation upon being grown to full confluence (Fig. 5a, Supplementary Fig. 6a). By contrast, HEK293A cells depleted of STK25 exhibited reduced levels of YAP phosphorylation as assessed by quantitative immunoblotting (Fig. 5a) and phos-tag analysis (Supplementary Fig. 6b). We also observed reduced TAZ phosphorylation and subsequent stabilization of unphosphorylated TAZ levels following loss of STK25 (Supplementary Fig. 6c). These results were reproduced in STK25 KO 293A cells (Supplementary Fig. 6d). We also grew adherent cells in suspension for defined periods of time, as cell detachment is another known activator of LATS activity (Zhao et al. 2012). We found that depletion of STK25 significantly impairs the dynamics of YAP phosphorylation under these conditions, such that not only does the overall magnitude of YAP phosphorylation become blunted, but YAP gets phosphorylated more slowly and to a lesser extent in STK25-depleted cells compared to controls (Fig. 5c). Together, these results demonstrate that STK25 plays a significant role in activating LATS kinases following cellular perturbations known to activate the Hippo signaling pathway.

**Figure 5.**
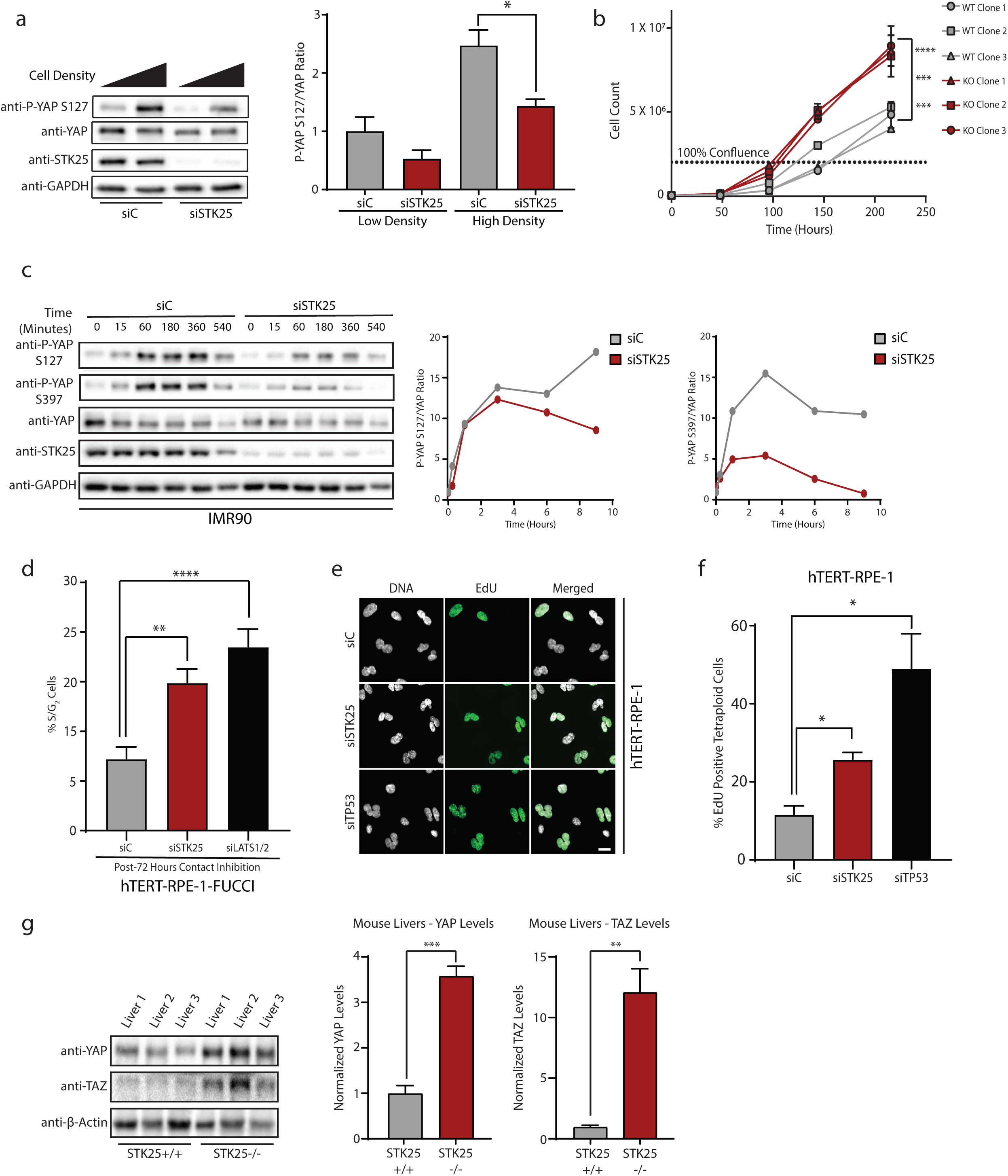
STK25 regulates YAP phosphorylation in response to physiologic stimuli. **a.** Immunoblot and quantification of YAP phosphorylation in HEK293A cells grown to low confluence or high confluence after transfection with the indicated siRNA. (n=3; *p<0.05, unpaired t-test). **b.** Cellular proliferation curves of control HEK293A clones and STK25 KO clones over the indicated time periods (n=3 replicates per cell line; ***p<0.001, ****p<0.0001, two-way ANOVA with Tukey’s post-hoc test). **c.** Representative immunoblot and quantification of YAP phosphorylation in IMR90 fibroblasts transfected with the indicated siRNA and held in suspension for the indicated time periods. **d.** Quantification of the percentage of cells remaining in S/G_2_ phase following prolonged contact inhibition in hTERT-RPE-1-FUCCI cells transfected with the indicated siRNA (n=4; **p<0.01, ****p<0.0001, one-way ANOVA with Dunnett’s post-hoc test). **e.** Cytokinesis failure was pharmacologically induced in hTERT-RPE-1 cells to generate binucleated tetraploid cells, and the percentage of EdU positive tetraploid cells following siRNA transfection was quantified. Cells were stained for DNA (White) and EdU incorporation (Green). Scale bar, 20 µm. **f.** Quantification of the percentage of EdU positive binucleated tetraploid cells following transfection with the indicated siRNA. TP53 siRNA served as positive control. (n=4 biological replicates; *p<0.05, One-way ANOVA with Dunnett’s post-hoc test). **g.** Protein samples were collected from livers of *STK25*+/+ and *STK25*-/- adult mice. Immunoblot and quantification of YAP and TAZ levels in these mice are presented (n=3 per genotype, **p<0.01, ***p<0.001, unpaired t-test). All data is presented as mean ± SEM unless otherwise indicated.

We hypothesized that depletion of STK25, with subsequent YAP activation, would also provide proliferative advantages to cells. Indeed, we found that cells genetically depleted of STK25 have an increased growth rate in culture (Fig. 5b). We also found that depletion of STK25 allows cells to partially overcome cell cycle arrest induced by contact inhibition, thereby validating our findings that loss of STK25 prevents YAP phosphorylation under contact inhibited conditions (Fig. 5d). Further, it has previously been demonstrated that tetraploid cells, which arise from cytokinetic failures, fail to proliferate efficiently due to LATS activation and subsequent inactivation of YAP. Proliferation can be restored to tetraploid cells through restoration of YAP activity (Ganem et al. 2014). Indeed, we found that STK25 knockdown is also sufficient to restore proliferative capacity to tetraploid cells (Fig. 5e, f). Lastly, we wished to assess if loss of STK25 would have similar effects *in vivo*. To do this, we extracted protein samples from the livers of *STK25*-/- adult mice and their wild-type adult littermates, which we then analyzed via SDS-PAGE for YAP and TAZ levels (Fig. 5g). As expected, we found increased levels of YAP and TAZ protein in the *STK25*-/- mice livers compared to their wild-type controls, indicating that loss of STK25 inactivates Hippo signaling under physiologic conditions both *in vitro* and *in vivo* (Fig. 5g).

Our data suggest STK25 is a novel activator of the LATS kinases whose depletion leads to activation of YAP/TAZ and subsequent cellular proliferation. YAP/TAZ are known to be hyper-activated across a wide range of human malignancies, although the mechanisms leading to their activation remain poorly understood. Our data suggest that loss of STK25 may represent one route through which cancer cells functionally activate YAP/TAZ. Interestingly, bioinformatic analysis of the TCGA reveals that focal deletion of *STK25* is a very common event across many tumor subtypes, with deep deletions occurring in a significant proportion of multiple aggressive cancers (Fig. 6a, b). For example, *STK25* is homozygously deleted from nearly 9% of all sarcomas (Fig 6a, b). We assessed if *STK25* deletion status has an effect on the clinical course of disease and found that sarcoma patients with *STK25* deletions have significantly shorter durations of survival than those with intact *STK25* and that deletions of *STK25* are more common in patients with recurrent disease than those without (Fig. 6c, Supplementary Fig. 7a). Additionally, we found that this pattern of recurrent focal loss in human cancers is unique to *STK25* among the currently identified LATS activating kinases (Supplementary Fig. 7b). Taken together, our findings suggest that loss of *STK25* may be one mechanism through which human cancers functionally inactivate Hippo signaling to promote tumorigenesis and disease progression.

**Figure 6.**
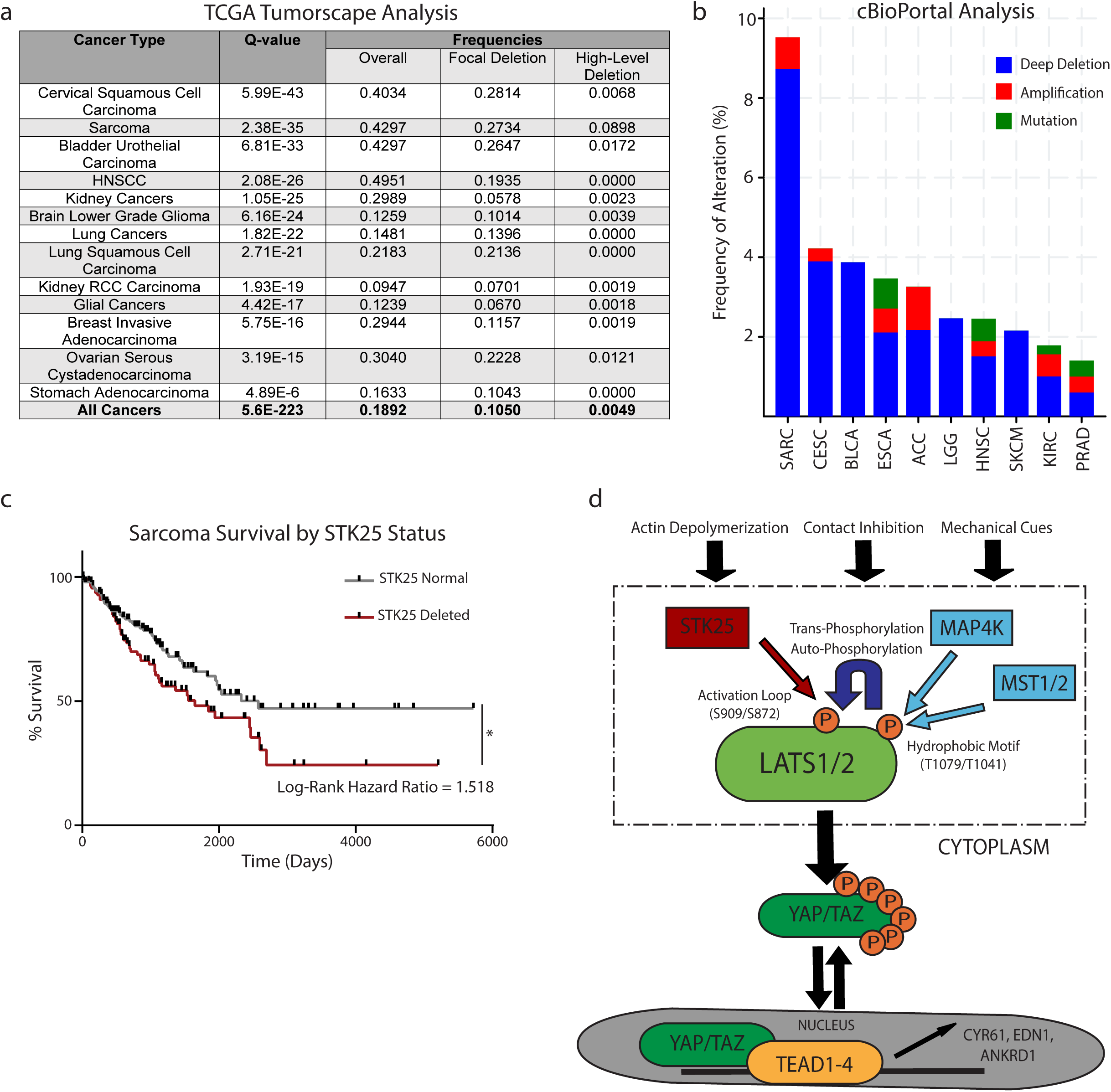
Loss of STK25 is common in human cancers and adversely affects patient survival. **a.** Publicly available TCGA datasets were probed to assess rates of focal deletion in *STK25* using the Tumorscape program online (http://www.broadinstitute.org/tcga/). The top 10 cancers with the highest rates of focal deletions in *STK25* are shown, together with the “All Cancers” dataset. **b.** Graphical representation of human cancers with the highest frequencies of *STK25* deletion. Data was accessed using the cBioPortal online program (http://www.cbioportal.org/). **c.** Survival data of sarcoma patients from the TCGA dataset were accessed using the Xenabrowser online program (https://xenabrowser.net/) and overall survival rates and times were assessed for patients with and without deletions of *STK25* (*p=0.0172, n=215, log-rank test). **d.** Proposed model of STK25 in Hippo tumor suppressor signaling.

## Discussion

It has been well established since the first knockout studies that deletions of both *MST1* and *MST2* are not sufficient to completely inactivate LATS signaling in mammalian cells (Zhou et al. 2009), strongly indicating the presence of additional LATS regulators (Thompson and Sahai 2015). Indeed, members of the MAP4K family, including human orthologs of *Drosophila* Hppy (MAP4K1, 2, 3, and 5) and Msn (MAP4K4, 6, and 7) were recently identified as upstream activators of LATS kinases. Like MST1/2, MAP4K proteins activate LATS via phosphorylation of the LATS hydrophobic motif (Li et al. 2014, Zheng et al. 2015, Meng et al. 2015, Hoa et al. 2016). The presence of such redundancies highlights the critical nature of LATS signaling in ensuring that cellular proliferation ceases when appropriate. In contrast, STK25-dependent activation of LATS entirely bypasses the need for phosphorylation at the hydrophobic motif, which provides some explanation for the robustness with which loss of this singular kinase is able to blunt LATS activation. Further, our study has revealed that LATS phosphorylation at the activation loop is not solely an auto- or trans-phosphorylation event, as we have identified STK25 as the first kinase able to phosphorylate the LATS activation loop motif independent of LATS kinase activity. It has previously been shown by several groups that while genetic deletion of the MST/MAP4K pathway components potently decreases levels of LATS and YAP phosphorylation, it is still possible to induce additional LATS activity upon treatment with actin depolymerizing agents or contact inhibition (Zhang et al. 2015, Meng et al. 2015, Li et al. 2015, Plouffe et al. 2016). We have now shown that STK25 is responsible for at least a portion of this remaining LATS activity, and that the mechanism through which it promotes LATS activity is novel and independent from other upstream Hippo kinases. This may explain the robustness with which loss of this singular kinase is able to potently activate YAP. Further, this lack of mechanistic redundancy may also explain why STK25 appears to be frequently focally deleted in a spectrum of human cancers, which is not observed with members of the MST/MAP4K signaling cascade (Supplementary Fig. 7b).

Several questions remain regarding the regulation of Hippo signaling, especially with respect to how stimulatory inputs that induce loss of cytoskeletal tension ultimately activate kinases upstream of LATS (Zhou et al. 2009, Zheng et al. 2015, Meng et al. 2015). While the activation of MAP4K proteins is poorly understood, MST1/2 are activated by both the TAO family kinases and through auto-phosphorylation events (Praskova et al. 2004, Boggiano et al. 2011, Poon et al. 2011), suggesting STK25 may be subject to similar regulatory control. Alternatively, STK25 may be constitutively active and loss of cytoskeletal tension simply promotes LATS-STK25 interaction. Indeed, it was recently shown that TRIP6 negatively regulates LATS by competing for MOB1 binding, and that this inhibition of LATS is relieved upon loss of cytoskeletal tension, suggesting that loss of tension allows for recruitment of LATS to other binding partners that promote its activity (Dutta et al. 2018). Another possibility involves spatial regulation of STK25 in tandem with the status of the Golgi apparatus. STK25 is known to localize to and regulate the polarization of the Golgi ribbon (Preisinger et al. 2004, Matsuki et al. 2010). Recent studies have implicated the Golgi as serving a sensor role that integrates extracellular signals for signaling pathways regulating cellular proliferation (Thomas et al. 2014). It is possible that when the Golgi becomes disorganized under conditions of low cytoskeletal tension, such as when a cell becomes contact inhibited or actin becomes pharmacologically disrupted (Lazaro-Dieguez et al. 2006), that STK25 becomes released from the Golgi to associate with LATS kinases. Alternatively, loss of tension and consequent Golgi disruption may serve as a signal to recruit LATS kinases to sites of disturbed Golgi, where they can then associate with STK25 to promote LATS activity.

In conclusion, we define STK25 as a novel regulator of Hippo signaling, which activates LATS1/2 via a novel mechanism independent from the canonical MST/MAP4K signaling pathway. The novelty of this mechanism explains why loss of this single kinase is able to induce significant activation of YAP, which is unable to be compensated for by the presence of other upstream Hippo kinases. We posit that *STK25* is a putative tumor suppressor gene, with data from human cancers supporting our claim; *STK25* appears to undergo significant focal deletions in a large variety of human cancers, and loss of STK25 in our cellular models promotes increased proliferation and resistance to stimuli that would normally induce cell cycle arrest. Deletions or mutations of core Hippo pathway components are rare, and it remains to be discovered how cancer cells overcome this critical tumor suppressor pathway in the process of transformation. Our data demonstrate that loss of STK25 represents one potential route through which cancer cells might deregulate this critical tumor suppressor pathway to achieve pathologic capacity.

## Acknowledgments

We would like to thank Kun-Liang Guan and members of his lab for technical advice and cell lines, Duojia Pan for reagents, and Bob Varelas and Anurag Singh for critical discussions about the work. We would also like to thank members of the Ganem lab for detailed comments on the manuscript. S.L is supported by a Medical Student Research Grant from the Melanoma Research Foundation and a Medical Student Young Investigator Award from the American Skin Association. M.V. is supported by a training grant from the NIH/NIGMS (5T32GM008541-20) and an F30 Award from the NCI (1F30CA228388-01). R.Q. is supported by a Canadian Institutes of Health Research Doctoral Foreign Study Award. B.H. is supported by NIH grant NS073662. N.G is a member of the Shamim and Ashraf Dahod Breast Cancer Research Laboratories and is supported by NIH grants CA154531 and GM117150, the Karin Grunebaum Foundation, the Smith Family Awards Program, the Melanoma Research Alliance, and the Searle Scholars Program.

## Author Contributions

S.L., H.C., and N.J.G. conceptualized and designed the study. S.L. performed experiments with R.J.Q. for bioinformatic analysis of datasets and with T.M., H.M.M., M.A.V., and I.P.M.M. for all other experiments. S.L. and N.J.G. analyzed the results. B.W.H. provided critical reagents. S.L. and N.J.G. wrote the manuscript. All authors commented on the manuscript.

## References

1. Adler, J.J. et al. 2013. Serum deprivation inhibits the transcriptional co-activator YAP and cell growth via phosphorylation of the 130-kDa isoform of Angiomotin by the LATS1/2 protein kinases. Proc Natl Acad Sci. 110:17368-73.

2. Boggiano, J.C., Vanderzalm, P.J., & Fehon, R.G. 2011. Tao-1 phosphorylates Hippo/MST kinases to regulate the Hippo-Salvador-Warts tumor suppressor pathway. Dev Cell. 21: 888-895.

3. Dupont, S. et al. 2011. Role of YAP/TAZ in mechanotransduction. Nature. 474:179-183.

4. Dutta, S. et al. 2018. TRIP6 inhibits Hippo signaling in response to tension at adherens junctions. EMBO Rep. 19: 337-350.

5. Enzo, E. et al. 2015. Aerobic glycolysis tunes YAP/TAZ transcriptional activity. EMBO J. 34: 1349-1370.

6. Fernandez-L, A. et al. 2009. YAP1 is amplified and up-regulated in hedgehog-assisted medulloblastomas and mediates Sonic hedgehog-driven neural precursor proliferation. Genes Dev. 23: 2729-2741.

7. Ganem, N.J. et al. 2014. Cytokinesis failure triggers hippo tumor suppressor pathway activation. Cell. 158:833-848.

8. Hoa, L. et al. 2016. The characterisation of LATS2 kinase regulation in Hippo-YAP signalling. Cell Signal. 28: 488-497.

9. Hong, J.H. et al. 2004. TAZ, a transcriptional modulator of mesenchymal stem cell differentiation. Science. 309:1074-1078.

10. Kelliher, F.C. & O’Sullivan, H. 2013. Oxford and the Savannah: Can the Hippo Provide and Explanation for Peto’s Paradox? Clin Cancer Res. 20:1-8.

11. Kim, M.H. et al. 2016. Actin remodeling confers BRAF inhibitor resistance to melanoma cells through YAP/TAZ activation. EMBO J. 35:462-78.

12. Lazaro-Dieguez, F. et al. 2006. Actin filaments are involved in the maintenance of Golgi cisternae morphology and intra-Golgi pH. Cell Motil Cytoskeleton. 63: 778-791.

13. Li, Q. et al. 2014. The conserved misshapen-warts-Yorkie pathway acts in enteroblasts to regulate intestinal stem cells in Drosophila. Dev. Cell. 31: 291-304.

14. Li, S., Cho, Y.S., Yue, T., Ip, Y.T., & Jiang, J. 2015. Overlapping functions of the MAP4K family kinases Hppy and Msn in Hippo signaling. Cell Discov. 1: 15038.

15. Li, W. et al. 2014. Merlin/*NF2* Loss-Driven Tumorigenesis Linked to CRL4^DCAF1^-Mediated Inhibition of the Hippo Pathway Kinases Lats1 and 2 in the Nucleus. Cancer Cell. 14: 48-60.

16. MacLean-Fletcher, S. & Pollard, T.D. 1980. Mechanism of action of cytochalasin B on actin. Cell. 20:329-341.

17. Matsuki, T. et al. 2010. Reelin and stk25 have opposing roles in neuronal polarization and dendritic Golgi deployment. Cell. 143:826-836.

18. Meng, Z., Moroishi, T., & Guan, K.L. 2016. Mechanisms of Hippo pathway regulation. Genes Dev. 30:1-17.

19. Meng, Z. et al. 2015. MAP4K family kinases act in parallel to MST1/2 to activate LATS1/2 in the Hippo pathway. Nat Commun. 6:8357.

20. Mo, J.S. et al. 2015. Cellular energy stress induces AMPK-mediated regulation of YAP and the Hippo pathway. Nat Cell Biol. 17:500-510.

21. Mohseni, M. et al. 2014. A genetic screen identifies an LKB1-MARK signaling axis controlling the Hippo-YAP pathway. Nat Cell Biol. 16: 108-117.

22. Mori, M. et al. 2014. Hippo signaling regulates microprocessor and links cell-density dependent miRNA biogenesis to cancer. Cell 156: 893-906.

23. Morton, W.M., Ayscough, K.R., & McLaughlin, P.J. 2000. Latrunculin alters the actin-monomer subunit interface to prevent polymerization. Nat Cell Biol. 2: 376-378.

24. Ni, L., Zheng, Y., Hara, M., Pan, D., & Luo, X. 2015. Structural basis for Mob1-dependent activation of the core Mst-Lats kinase cascade in Hippo signaling. Genes Dev. 29: 1416-1431.

25. Nishio, M. et al. 2016. Dysregulated YAP1/TAZ and TGF-β signaling mediate hepatocarcinogenesis in *Mob1a/1b*-deficient mice. Proc Natl Acad Sci. 113:E71-E80.

26. Overholtzer, M. et al. 2006. Transforming properties of YAP, a candidate oncogene on the chromosome 11q22 amplicon. Proc Natl Acad Sci. 103:12405-12410.

27. Pan, D. 2010. The hippo signaling pathway in development and cancer. Dev Cell. 19: 491-505.

28. Park, Y.Y. et al. 2016. YAP1 and TAZ Activates mTORC1 Pathway by Regulating Amino Acid Transporters in hepatocellular carcinoma. Hepatology 63: 159-172.

29. Plouffe, S.W. et al. 2016. Characterization of Hippo Pathway Components by Gene Inactivation. Mol Cell 64:993-1008.

30. Poon, C.L., Lin, J.I., Zhang, X., & Harvey, K.F. 2011. The sterile 20-like kinase Tao-1 controls tissue growth by regulating the Salvador-Warts-Hippo pathway. Dev Cell. 21: 896-906.

31. Praskova, M., Khoklatchev, A., Ortiz-Vega, S., & Avruch, J. 2004. Regulation of the MST1 kinase by autophosphorylation, by the growth inhibitory proteins, RASSF1 and NORE1, and by Ras. Biochem J. 381: 453-462.

32. Preisinger, C. et al. 2004. YSK1 is activated by the Golgi matrix protein GM130 and plays a role in cell migration through its substrate 14-3-3zeta. J Cell Biol. 164:1009-1020.

33. Stegert, M.R., Tamaskovic, R., Bichsel, S.J., Hergovich, A., & Hemmings, B.A. 2004. Regulation of NDR2 Protein Kinase by Multi-site Phosphorylation and the S100B Calcium-binding Protein. J Biol Chem. 279: 23806-23812.

34. Tamaskovic, R., Bichsel, S.J., Rogniaux, J., Stegert, M.R., & Hemmings, B.A. 2003. Mechanism of Ca^2+^-mediated Regulation of NDR Protein Kinase through Autophosphorylation and Phosphorylation by an Upstream kinase. J Biol Chem. 278: 6710-6718.

35. Thomas, J.D. et al. 2014. Rab1A is an mTORC1 Activator and Colorectal Oncogene. Cancer Cell. 26: 754-769.

36. Thompson, B.J., & Sahai, E. 2015. MST kinases in development and disease. J Cell Biol. 210:871-82.

37. Wang, W. et al. 2015. AMPK modulates Hippo pathway activity to regulate energy homeostasis. Nat Cell Biol. 17:490-499.

38. Wu, S., Liu, Y., Zheng, Y., Dong, J., & Pan, D. 2008. The TEAD/TEF family protein Scalloped mediates transcriptional output of the Hippo growth-regulatory pathway. Dev Cell. 14:388-98.

39. Yin, F. et al. 2013. Spatial Organization of Hippo signaling at the plasma membrane mediated by the tumor suppressor Merlin/NF2. Cell. 154:1342-1355.

40. Yu, F.X. et al. 2012. Regulation of the Hippo-YAP pathway by G-protein-coupled receptor signaling. Cell. 150:780-791.

41. Zanconato, F., Cordenonsi, M., & Piccolo, S. 2016. YAP/TAZ at the Roots of Cancer. Cancer Cell. 29:783-803.

42. Zanconato, F. et al. 2015. Genome-wide association between YAP/TAZ/TEAD and AP-1 at enhancers drives oncogenic growth. Nat Cell Biol. 17:1218-1227.

43. Zhang, H. et al. 2009. TEAD transcription factors mediate the function of TAZ in cell growth and epithelial-mesenchymal transition. J Biol Chem. 284:13355-13362.

44. Zhang, L., Ren, F., Zhang, Q., Chen, Y., Wang, B., & Jiang, J. 2008. The TEAD/TEF family of transcription factor Scalloped mediates Hippo signaling in organ size control. Dev Cell. 14:377-87.

45. Zhao, B., Li, L., Tumaneng, K., Wang, C.Y., & Guan, K.L. 2010. A coordinated phosphorylation by Lats and CK1 regulates YAP stability through SCF(beta-TRCP). Genes Dev. 24:72-85.

46. Zhao, B. et al. 2012. Cell detachment activates the Hippo pathway via cytoskeleton reorganization to induce anoikis. Genes Dev. 26:54-68.

47. Zhao, B. et al. 2007. Inactivation of YAP oncoprotein by the Hippo pathway is involved in cell contact inhibition and tissue growth control. Genes Dev. 21:2747-2761.

48. Zheng, Y. et al. 2015. Identification of Happyhour/MAP4K as Alternative Hpo/Mst-like Kinases in the Hippo Kinase Cascade. Dev Cell. 34:642-55.

49. Zhou, D. et al. 2009. Mst1 and Mst2 maintain hepatocyte quiescence and suppress hepatocellular carcinoma development through inactivation of the Yap1 oncogene. Cancer Cell. 19: 425-438.

