## Supplementary Materials for "STK25 directly activates LATS1/2 independent of MST/MAP4Ks"

Supplementary Figure 1

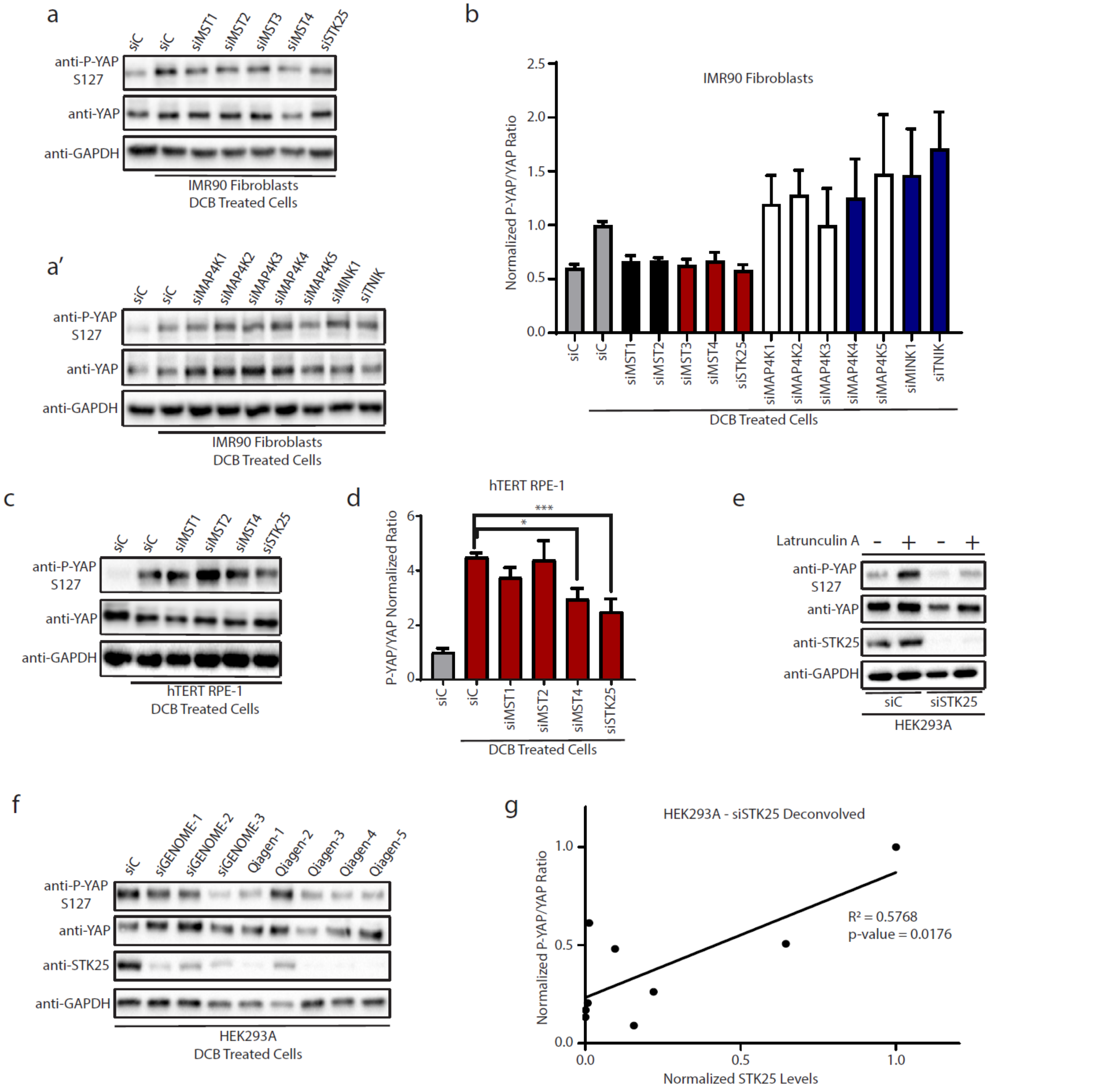


**Supplementary Figure 1. Loss of STK25 decreases YAP phosphorylation in response to actin disruption.** **a.** Representative immunoblot of YAP phosphorylation following treatment with 10 µM DCB in IMR90 fibroblasts transfected with the indicated siRNA. **b.** Quantitation of the IMR90 focused kinome screen (n=3, *$p<0.05$ by One-Way ANOVA with Dunnett's post-hoc analysis). **c.** Representative immunoblot of YAP phosphorylation following treatment with 10 µM DCB in hTERT-RPE-1 cells transfected with the siRNA. **d.** Quantitation of the RPE-1 secondary kinome screen (n=4, *$p<0.05$, ***$p<0.001$ by One-Way ANOVA with Dunnett's post-hoc analysis). **e.** Representative immunoblot of YAP phosphorylation following treatment with 1 µg/mL Latrunculin A in HEK293A cells transfected with the indicated siRNA. **f.** Immunoblot and **g.** quantification of YAP phosphorylation and STK25 protein levels following treatment with 10 µM DCB in HEK293A cells transfected with the indicated siRNA. STK25 protein levels were normalized to GAPDH protein levels and plotted against the respective P-YAP/YAP ratios. Linear regression was performed to assess whether a correlation existed between levels of STK25 protein expression and P-YAP/YAP ratios (p=0.0176, Pearson's $R^2 = 0.5768$).

Supplementary Figure 2

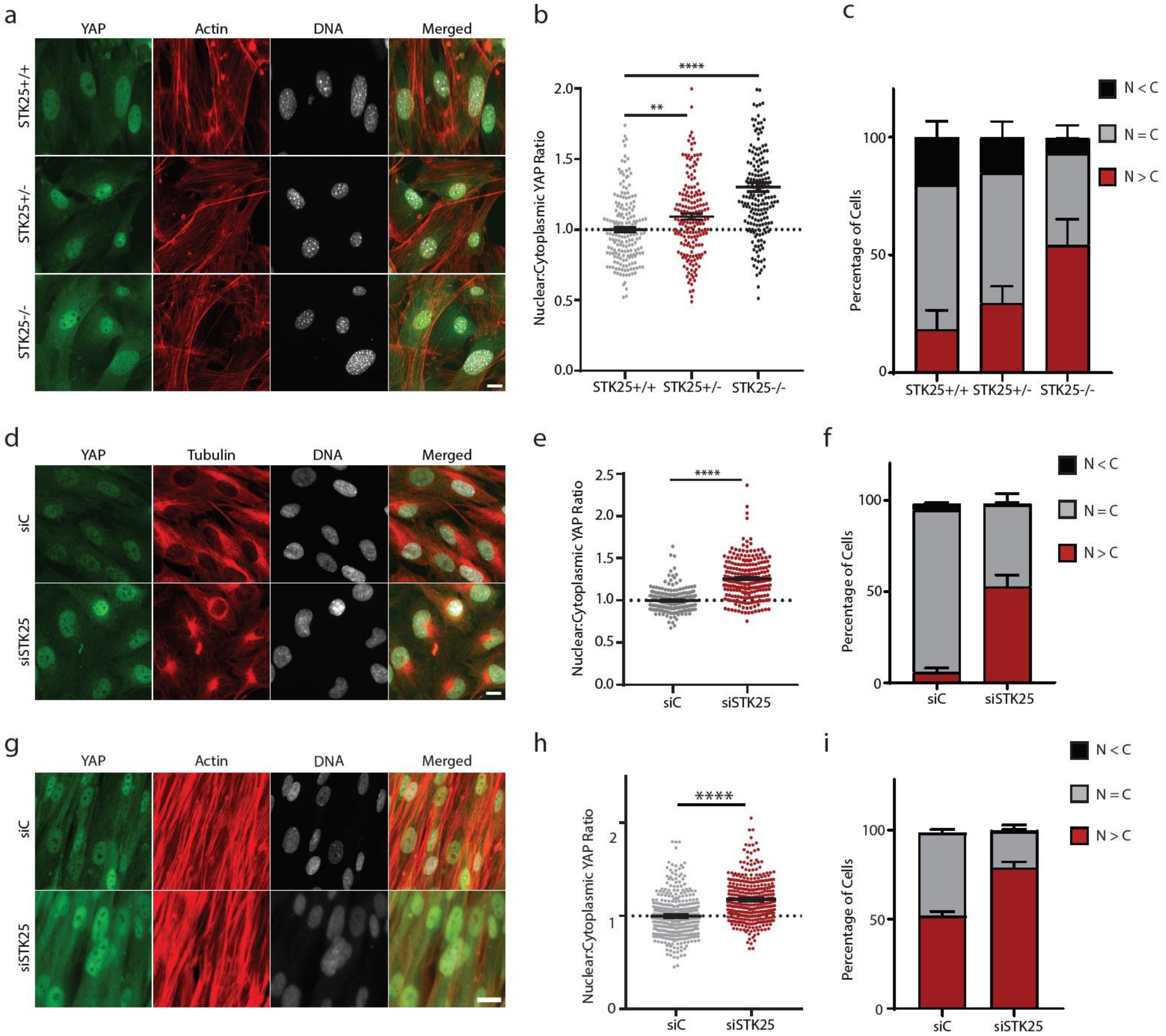


**Supplementary Figure 2. Loss of STK25 increases levels of active, nuclear YAP.**
**a.** MEFs isolated from STK25+/+, STK25+/-, and STK25-/- mice were plated on coverslips and stained for YAP (Green), Actin (Red), and DNA (White). Scale bar, 20 µm. **b.** YAP intensity was quantified and nuclear:cytoplasmic ratios were calculated for the MEF images (n>150 per group over 3 biological replicates; **p<0.01, ****p<0.0001 by Kruskal-Wallis test with Dunn's post-test). **c.** YAP localization in the MEFs was quantified (n=3 biological replicates, N>C, YAP is enriched in the nucleus; N=C, YAP is evenly distributed between the nucleus and the cytoplasm; N<C, YAP is enriched in the cytoplasm). **d.** hTERT-RPE-1 transfected with the indicated siRNA were stained for YAP (Green), Tubulin (Red), and DNA (White). Scale bar, 20 µm. **e.** YAP intensity was quantified and nuclear:cytoplasmic ratios were calculated for the RPE-1 experiments (n=225 over 3 biological replicates; ****p<0.0001 by Mann-Whitney test). **f.** YAP localization in the hTERT-RPE-1 cells was quantified (n=3 biological replicates, N>C, YAP is enriched in the nucleus; N=C, YAP is evenly distributed between the nucleus and the cytoplasm; N<C, YAP is enriched in the cytoplasm). **g.** IMR90 fibroblasts transfected with the indicated siRNA were stained for YAP (Green), Actin (Red), and DNA (White). Scale bar, 20 µm. **h.** YAP intensity was quantified and nuclear:cytoplasmic ratios were calculated (n=300 per group over 4 biological replicates; ****p<0.0001, Mann-Whitney test). **i.** YAP localization in the IMR90 cells was quantified (n=4 biological replicates, N>C, YAP is enriched in the nucleus; N=C, YAP is evenly distributed between the nucleus and the cytoplasm; N<C, YAP is enriched in the cytoplasm).

Supplementary Figure 3

a

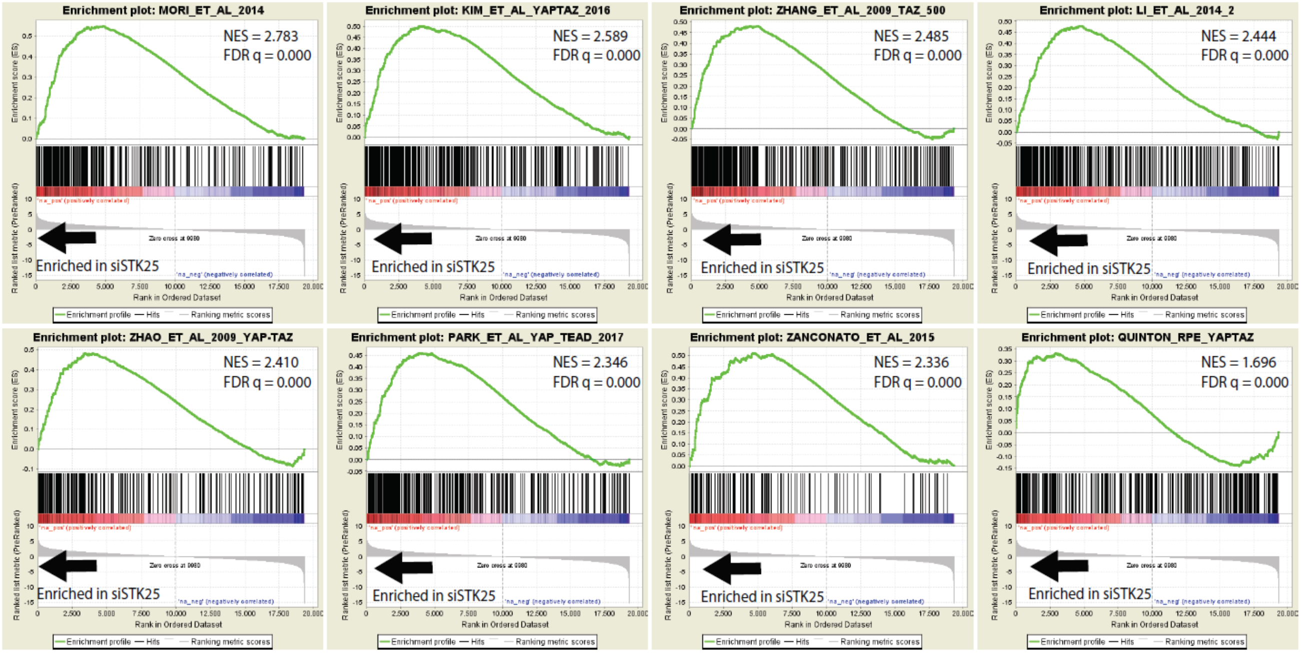


b

| Annotated Cellular Function/ Published YAP/TAZ gene set | Enrichment Score | Normalized Enrichment Score | FDR q-value |
|---|---|---|---|
| **Park et al. 2016** | **0.62** | **3.17** | **0.000** |
| *HALLMARK_MYC_TARGETS_V1* | *0.64* | *3.06* | *0.000* |
| *HALLMARK_E2F_TARGETS* | *0.63* | *3.04* | *0.000* |
| **Enzo et al. 2015** | **0.59** | **3.01** | **0.000** |
| **Mohseni et al. 2014** | **0.55** | **2.84** | **0.000** |
| **Mori et al. 2014** | **0.55** | **2.77** | **0.000** |
| *HALLMARK_OXIDATIVE_PHOSPHORYLATION* | *0.56* | *2.66* | *0.000* |
| **Kim et al. 2016 – YAP/TAZ** | **0.50** | **2.60** | **0.000** |
| **Zhang et al. 2009 - TAZ** | **0.48** | **2.50** | **0.000** |
| *HALLMARK_G2M_CHECKPOINT* | *0.51* | *2.46* | *0.000* |

**Supplementary Figure 3. Cells depleted of STK25 exhibit enrichment of active YAP/TAZ gene signatures. a.** A list of genes upregulated upon knockdown of STK25 were assessed by GSEA using a list of curated publicly available active YAP/TAZ gene sets. The remainder of the curated list, excluding the top three gene sets already presented, are shown here; with the exception of the Park et al. YAP-TEAD 2017 gene set, which can be found under the accession code GSE32597, and the Quinton YAP-TAZ gene set, which can be found under the accession code GSEXXXXX, the names of the gene sets correspond to the publications from which they were derived. **b.** A table showing relative enrichment of active YAP/TAZ gene sets in comparison to enrichment of Hallmark gene sets from the Molecular Signatures Database is presented.

Supplementary Figure 4

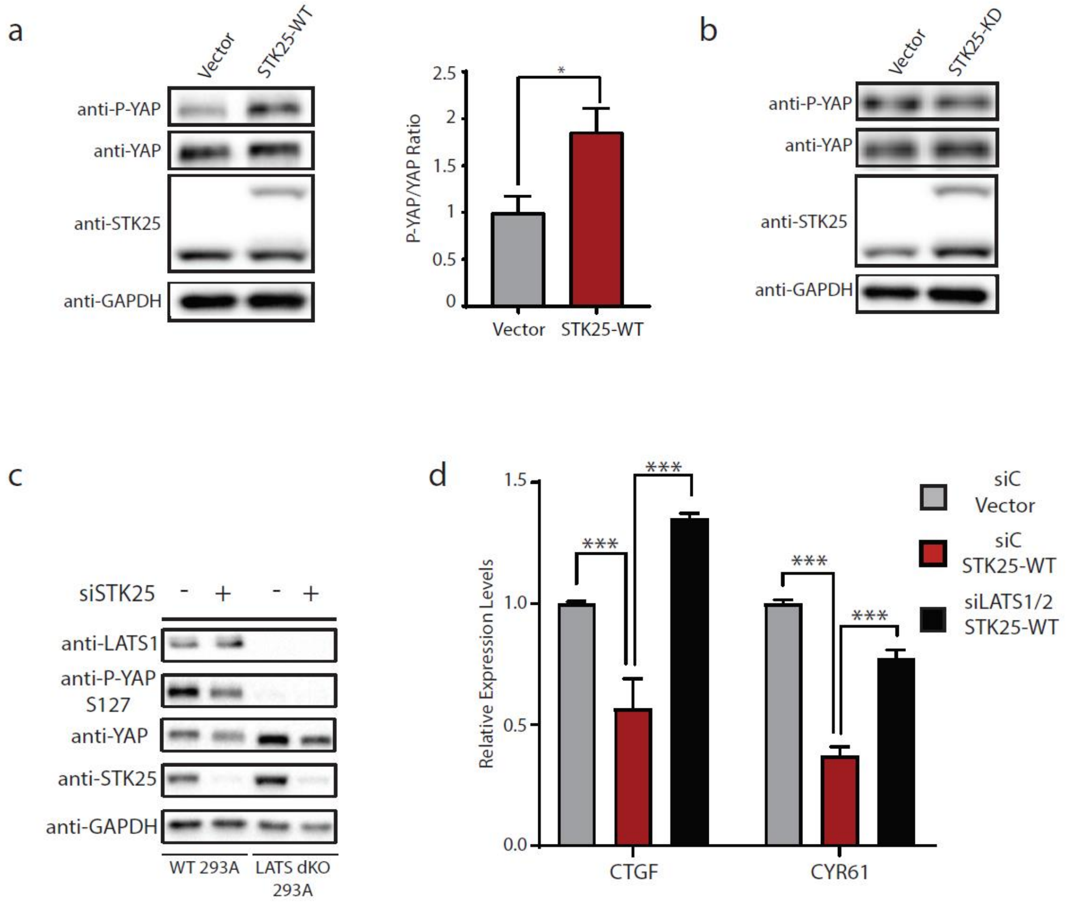


**Supplementary Figure 4. STK25 requires LATS1/2 for its inhibitory effects on YAP.**
**a.** Immunoblot and quantitation of phosphorylated YAP levels in HEK293A cells stably expressing wild-type STK25 (STK25-WT) or vector control (Vector) (n=3; *p<0.05, paired t-test). **b.** Immunoblot and quantitation of phosphorylation YAP levels in HEK293A cells stably expressing kinase-dead STK25 (STK25-KD) or vector control (Vector) (n=3; N.S. stands for not significant.) **c.** Representative immunoblot of phosphorylated YAP following STK25 knockdown in wild-type and LATS dKO HEK293A. Cells were grown to confluence in order to activate Hippo signaling to induce YAP phosphorylation. **d.** qRT-PCR analysis of YAP-target gene expression in HEK293A cells stably overexpressing either wild-type STK25 (STK25-WT) or vector control (Vector) after transfection with control siRNA or siRNAs targeting LATS1 and LATS2 (n=3; ***p<0.001, One-way ANOVA with Dunnett’s post-hoc analysis; DF=2, 6; F=30.08).

Supplementary Figure 5

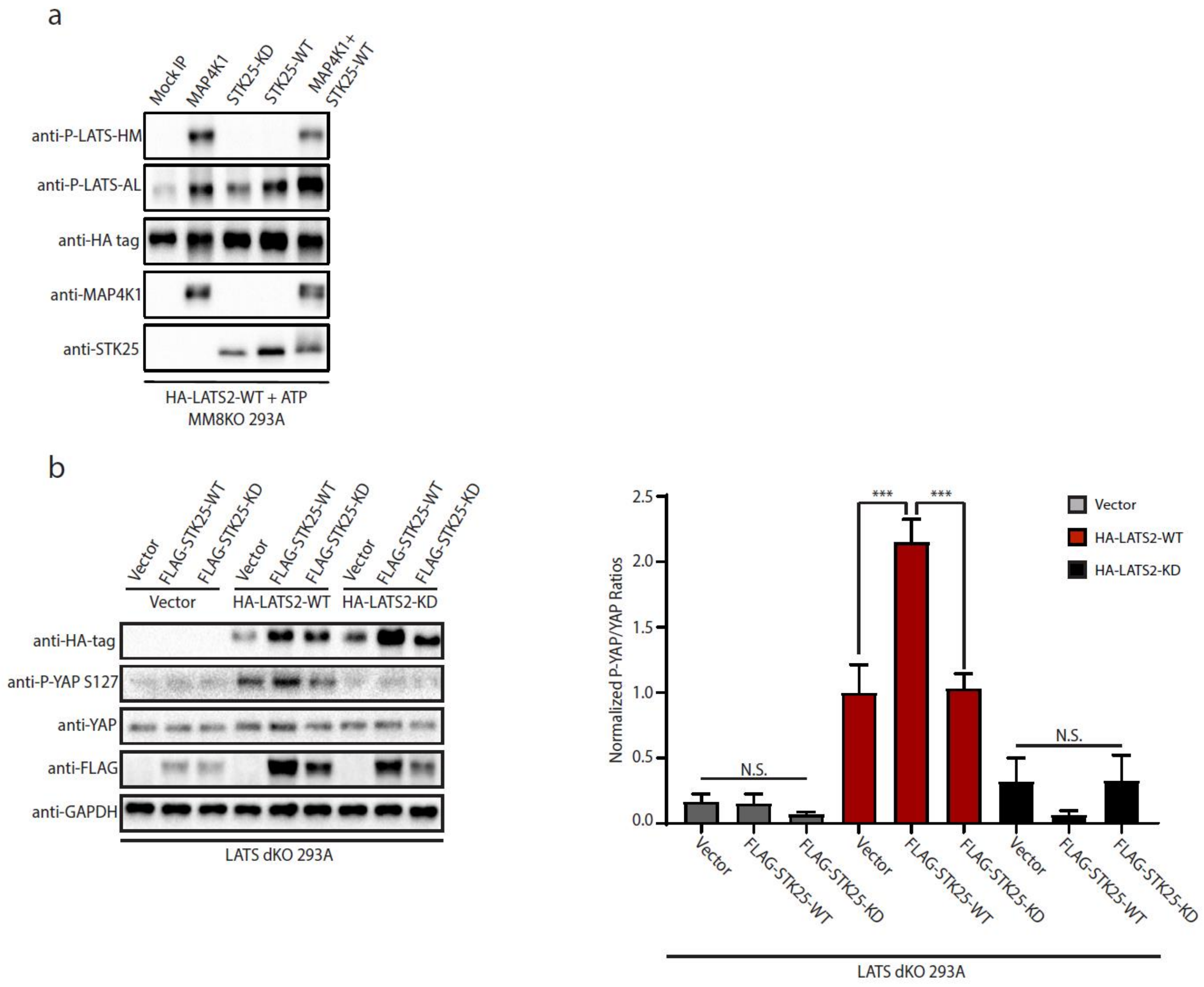


**Supplementary Figure 5. STK25 activates the LATS kinases.**
**a.** IP purified wild-type LATS2 (HA-LATS2-WT) from transfected MM8KO 293A cells was co-incubated with IP purified FLAG-STK25-WT, FLAG-STK25-KD, or FLAG-MAP4K1, all from transfected MM8KO 293A. Kinase reactions were allowed to occur in the presence of 500 μM ATP, and levels of phosphorylated LATS at the hydrophobic motif (P-LATS-HM) and activation loop motif (P-LATS-AL) were assessed via immunoblotting. A mock IP product from untransfected MM8KO 293A lysates using FLAG antibody and protein G magnetic beads served as control. **b.** LATS dKO 293A were transfected with empty vector, wild-type LATS2 (HA-LATS2-WT), or kinase-dead LATS2 (HA-LATS2-KD), alongside Vector, wild-type STK25 (FLAG-STK25-WT), or kinase-dead STK25 (FLAG-STK25-KD). Lysates were collected and assessed via SDS-PAGE and immunoblotting for YAP phosphorylation at serine127. Quantitation of YAP phosphorylation under the various transfection conditions are presented (n=3, ***p<0.001, One-Way ANOVA with Tukey's post-hoc test; DF=18, F=25.93).

Supplementary Figure 6

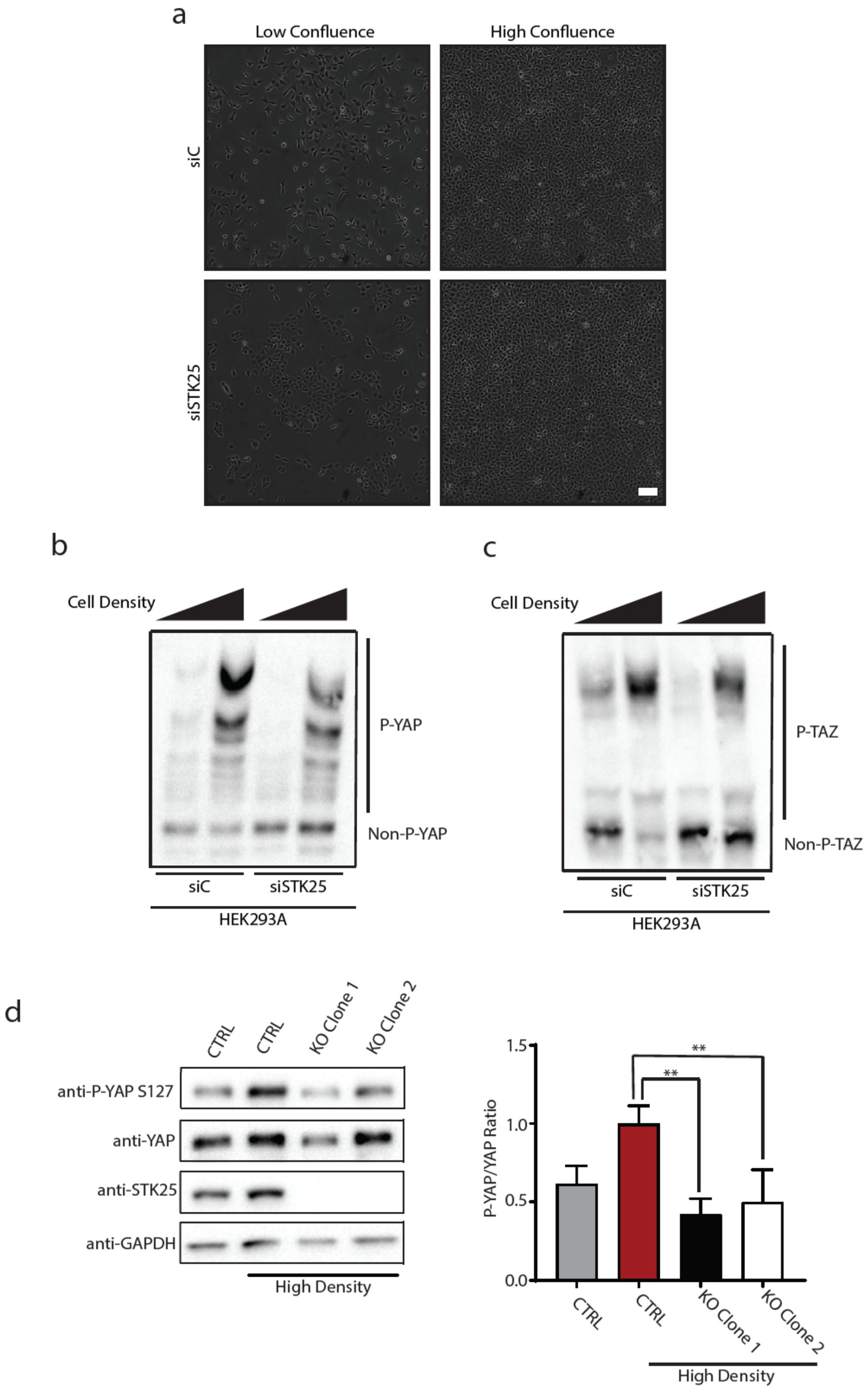


**Supplementary Figure 6. STK25 loss impairs physiologic Hippo activation.**
**a.** Representative phase images of HEK293A grown to low or high confluence following transfection with the indicated siRNA. Scale bar, 100 µm. **b.** Global phosphorylation status of YAP was assessed via phos-tag gel electrophoresis using lysates from HEK293A grown to either low or high confluence following transfection with the indicated siRNA. **c.** Global phosphorylation status of TAZ was assessed via phos-tag gel electrophoresis using lysates from HEK293A grown to either low or high confluence following transfection with the indicated siRNA. **d.** Immunoblot and quantitation of phosphorylated YAP levels under conditions of high confluence in either control HEK293A stably expressing Cas9 and a non-targeting sgRNA or STK25 KO 293A stably expressing Cas9 together with either sgRNA 1 (Clone 1) or sgRNA 2 (Clone 2) targeting STK25. (n=4; **p<0.01, One-way ANOVA with Dunnett's post-hoc analysis).

Supplementary Figure 7

a

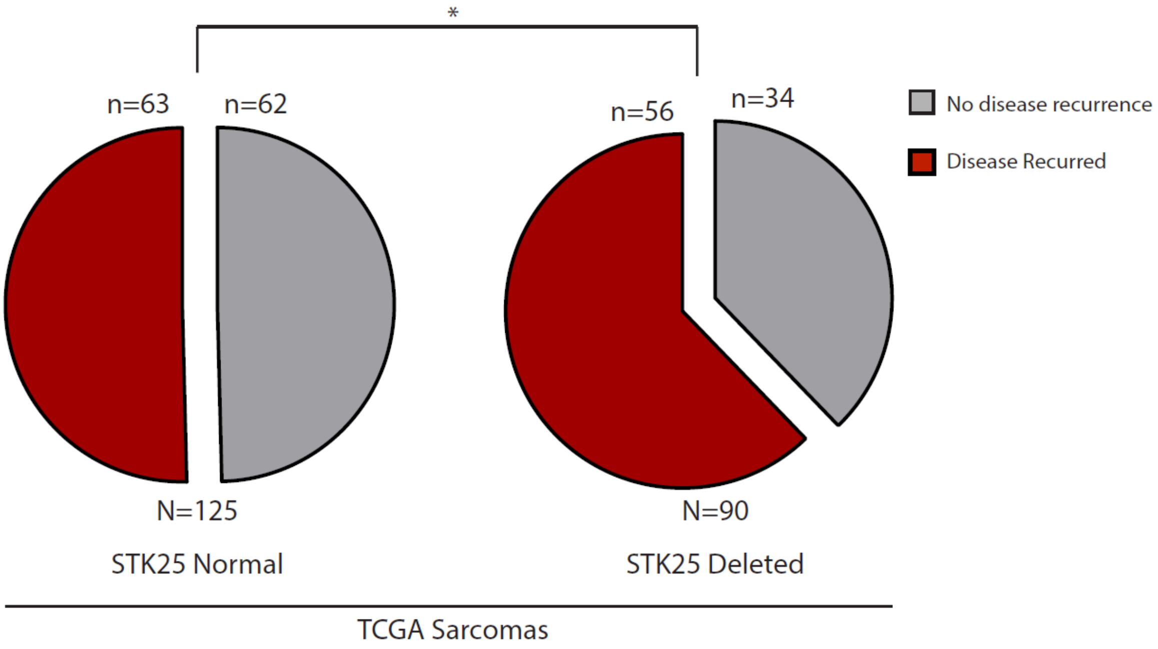


b

| Gene | Amplified/Deleted | Q-value | Overall Frequency |
|---|---|---|---|
| MST1 | Neither | N/A | N/A |
| **MST2** | **Amplified** | **1.07 E-15** | **0.4231** |
| **MAP4K1** | **Amplified** | **1.36 E-37** | **0.2158** |
| MAP4K2 | Neither | N/A | N/A |
| MAP4K3 | Neither | N/A | N/A |
| MAP4K4 | Neither | N/A | N/A |
| MAP4K5 | Neither | N/A | N/A |
| **TNIK/MAP4K6** | **Amplified** | **7.75 E-217** | **0.3284** |
| MINK1/MAP4K7 | Neither | N/A | N/A |
| **TAOK1** | **Amplified** | **1.55 E-7** | **0.1857** |
| TAOK2 | Neither | N/A | N/A |
| TAOK3 | Neither | N/A | N/A |
| **STK25** | **Deleted** | **5.61 E-223** | **0.1892** |

**Supplementary Figure 7. Focal deletions of identified Hippo pathway components are rare.**
**a.** Rates of recurrence-free survival in sarcoma patients with and without deletions in *STK25* (n=125 for patients without deletions, n=90 for patients with deletions, *$p<0.05$, two-tailed chi-square goodness-of-fit test).
**b.** Publicly available TCGA datasets were probed to assess rates of focal deletion in known upstream activators of LATS kinases, using the "All Cancers" dataset from the Tumorscape program online (http://www.broadinstitute.org/tcga/).
